# A prelimbic molecular clock of protein synthesis for memory persistence

**DOI:** 10.64898/2026.01.02.697403

**Authors:** J. Iqbal, S. Kim, S. Lawal, A. Shah, L. Punepalle, H. Sanghvi, A. Gallagher, A. Wilson, B. Xu, P Shrestha

## Abstract

Emotionally salient associative memories can endure for long periods, yet the mechanisms that determine their long-term stability remain unclear. Here we show that the prelimbic (PL) cortex integrates temporally structured translational programs to control both the consolidation and reconsolidation of cued threat memories. Using Pavlovian threat conditioning with *in vivo* fiber photometry, we found that PL calcium dynamics tightly track memory strength: discrete threat-predictive cues evoked robust activity during recent and single-timepoint remote retrieval, whereas prior retrieval selectively weakened remote expression, independent of contextual influences. Translational profiling of PL *Camk2a*⁺ cells uncovered a biphasic consolidation program, with an early phase characterized by ER stress-linked translational repression and robust oligodendrocyte plasticity, followed by a delayed phase engaging synaptic growth pathways. Loss- and gain-of-function approaches demonstrated that eIF2α-regulated, cap-independent translation is essential for recent consolidation and for the enduring stabilization of remote memory, whereas retrieval-induced destabilization engages a mechanistically distinct, eIF4E-dependent translational pathway required for reconsolidation. These findings identify the PL cortex as a dynamic node in which discrete modes of translational control govern the long-term persistence of emotional memories.

## Main

Emotionally salient associative memories can persist for extended period, sometimes even across the lifespan. Memory consolidation depends on tightly regulated, spatiotemporally precise protein synthesis across cortical and subcortical circuits^1–4^. Re-exposure to a unimodal cue previously paired with threat reactivates the underlying memory trace, eliciting robust, species-specific defensive responses^5,6^. Discrete threat-predictive cues generate far more precise and reliable responses than the multimodal contexts in which learning occurred^6^. Remote contextual-threat memories are thought to depend on enduring plasticity within the medial prefrontal cortex (mPFC), particularly the prelimbic (PL) cortex, which becomes progressively recruited as memory representations are redistributed from the medial temporal lobe to long-term cortical storage^7–9^. By contrast, unimodal cue-based threat memories rely minimally on hippocampal integrity^10,11^, raising the unresolved question of whether these discrete associations undergo analogous systems-level consolidation within the PL. Re-exposure to a threat-predictive cue also triggers retrieval-induced destabilization, briefly rendering the reactivated engram^12^ labile before it is re-stabilized through reconsolidation^3,13–16^. Yet, how a memory’s retrieval history shapes the durability and fidelity of remote threat memories - and whether this process is governed by translation-dependent mechanisms - remains poorly understood. Here, we define the prelimbic dynamics underlying memory encoding and retrieval, map temporally structured translational programs in the PL, and demonstrate a causal requirement for protein synthesis in both remote memory consolidation and retrieval-induced reconsolidation.

### Prelimbic cortex calcium dynamics track retrieval-dependent threat memory strength

Pavlovian cued-threat conditioning (PTC) provides a powerful paradigm for probing the neural mechanisms underlying emotional memory consolidation, as a single learning trial is sufficient to generate robust, long-lasting episodic memories thereby creating a clear boundary between learning and consolidation^17,18^. To investigate whether the PL activity is recruited during memory encoding and retrieval, we used *in vivo* fiber photometry in awake behaving mice to monitor pan-cellular calcium transients by virally expressing genetically encoded calcium indicator GCaMP6f^19^ and implanting optic ferrule within the PL (**Fig. 1a**). We opted to monitor calcium transients since they represent the movement of calcium ions (Ca²⁺) within and between cells and are essential for synaptic plasticity and the subsequent changes in gene expression that underpin memory storage^20,21^. We performed PTC by presenting mice with two pairings of an auditory tone (conditioned stimulus, CS) that co-terminated with a footshock (unconditioned stimulus, US) during training and tested remote long-term memory (LTM) for cued threat response in a novel context either at a single time-point (non-longitudinal, NL) or following recent retrieval in a longitudinal (L) framework (**Fig. 1b**). During the first CS-US pairing, we observed a large, rapid calcium transient in response to the US, which significantly decreased by 53% during the second pairing, indicating experience-dependent desensitization in the footshock-elicited PL response (**Fig. 1c, d**). On the other hand, this was accompanied with a 96% higher freezing response to the second CS that reflects behavioral learning (**Fig. 1e**).

**Figure 1.**
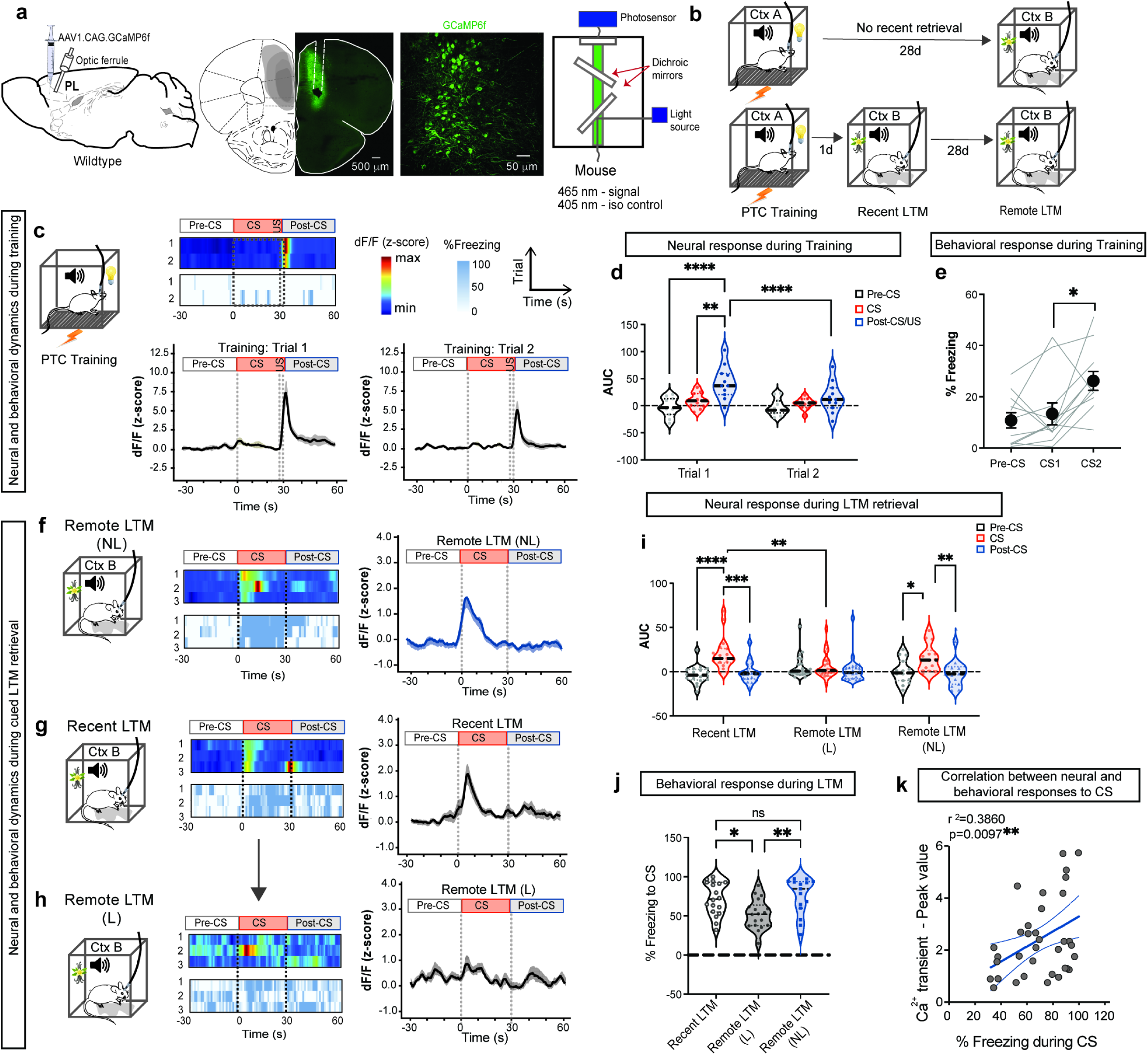
Prelimbic cortex calcium dynamics track retrieval-dependent threat memory strength. **a**) Schematic of *in vivo* fiber photometry showing AAV1.CAG.GCaMP6f injection and optic ferrule implant into the prelimbic (PL) cortex of wildtype mice to monitor neural activity via virally encoded GCaMP6f expression. Scale bars, 500 μm and 50 μm. **b)** Schematic of longitudinal and nonlongitudinal testing of remote LTM in Pavlovian threat conditioning (PTC) paradigm. Mice received two pairings of an auditory conditioned stimulus (CS) co-terminating with a foot shock (US) during training in Context A, followed by tests of recent and remote long-term memory (LTM) in Context B in longitudinal framework or that of single timepoint remote LTM at 28d post-training (non-longitudinal, NL). **c)** Representative heatmaps showing GCaMP6f fluorescence (ΔF/F) and freezing response during pre-CS, CS, and post-CS epochs (30s each) during training. Peri-event time histograms (PETH), mean+/-SEM, show that the first CS–US pairing elicited large PL calcium transients in response to the US, which decreased significantly during the second pairing, indicating experience-dependent attenuation of neural responses. **d)** Area-under-the-curve (AUC) analysis revealed significant differences between trials specifically during the post-CS/US period. Trial x CS Epoch: F (2, 12) = 4.335; CS Epoch: F (2, 12) = 8.274; n=5 mice/group. **e)** Freezing responses were significantly elevated during the second CS, reflecting behavioral learning. F (1.548, 15.48) = 8.549; n=11 animals/group. **f)** Representative heatmaps showing freezing response during pre-CS, CS, and post-CS epochs (30s each) during remote memory retrieval in the non-longitudinal (NL) group. PETH (mean+/-SEM) for ΔF/F z-scored traces demonstrate robust PL calcium activity during CS retrieval in the NL remote condition. **g, h)** Representative heatmaps showing freezing response during pre-CS, CS, and post-CS epochs (30s each) during recent and remote memory retrieval in the longitudinal (NL) group. PETH (mean+/-SEM) for ΔF/F traces demonstrate robust PL calcium activity during recent retrieval and retrieval-dependent modulation of remote memory. **i)** AUC analysis revealed significantly reduced CS-evoked PL activity during remote retrieval relative to recent memory in longitudinal conditions. CS Epoch X LTM: F (4, 162) = 3.841, CS Epoch: F (2, 162) = 18.43; n=5-8 mice/group. **j)** Freezing responses were significantly diminished at the remote timepoint relative to recent retrieval in longitudinal testing. F (2, 48) = 6.073; n=15-18 mice/group. **k)** Correlation analysis between CS-evoked calcium transients and freezing behavior during retrieval. r^2^=0.3860. **Statistical tests: d, i)** Two-way ANOVA with Bonferroni post-hoc test; **e)** RM One-way ANOVA with Bonferroni post-hoc test; **j)** One-way ANOVA with Bonferroni post-hoc test; **k)** Pearson correlation. *p<0.05, **p<0.01, ***p<0.001, ****p<0.0001, ns not significant.

During a single time-point NL-remote retrieval, CS evoked robust calcium transients in the prelimbic (PL) cortex, indicating prominent PL engagement (**Fig. 1f**). By contrast, the longitudinal group displayed strong CS-evoked calcium responses during recent LTM, which were markedly reduced during remote LTM (**Fig. 1g**, **h**). Consistent with this pattern, area-under-the-curve (AUC) analyses revealed robust CS-evoked activity above baseline during recent LTM and in the single-timepoint NL-remote LTM test. In contrast, during remote memory retrieval in the longitudinal group, CS responses did not differ from surrounding epochs and were reduced by 69% relative to recent LTM within the same subjects (**Fig. 1i**). The conditioned freezing responses paralleled the PL calcium dynamics: both recent retrieval and NL-remote retrieval were associated with high freezing levels, whereas L-remote retrieval showed a significant reduction (28%) in CS-elicited freezing (**Fig. 1j**). The inverse relationship between recent and remote memory strength could reflect context-dependent extinction–like processes arising from weakening of the tone–shock association following repeated CS exposure during recent retrieval. To directly test this possibility, we assessed remote LTM in a separate cohort of mice in a novel context (Context C). CS-evoked freezing during remote retrieval was indistinguishable between animals tested in Context B and Context C, indicating that context-dependent mechanisms do not account for the observed reduction in fear expression (**Extended Data Fig. 1a–b**). Moreover, this inverse coupling was specific to recent versus remote LTM, as neither behavioral performance nor PL neural responses were impaired at an intermediate retrieval time point (14 days post-learning; **Extended Data Fig. 2a–c**). Notably, the strong positive correlation between CS-evoked calcium transients in the PL and freezing behavior further supports the conclusion that PL activity provides a reliable neural readout of conditioned threat memory strength (**Fig. 1k**).

### Biphasic translational reprogramming during memory consolidation

At 15 minutes post-training, we observed rapid dephosphorylation of eIF2α in the prelimbic cortex (**Extended Data Fig. 3**), consistent with the surge in protein synthesis previously reported in the amygdala following Pavlovian threat conditioning. Building on this finding and having established that PL calcium activity is robustly engaged during both memory encoding and retrieval, we next examined how the translational landscape of calcium-responsive, *Camk2a*-expressing cells in the PL is dynamically regulated across the consolidation window at 1 hour and 4 hours post-training. *Camk2a* gene encodes a subunit of calcium/calmodulin-dependent protein kinase II, a multifunctional serine/threonine kinase that contributes to synaptic plasticity, largely via modulation of ionotropic glutamate receptors on dendritic spines and at postsynaptic densities, which in turn can influence synaptic strength^22^. CaMK2α typically assembles into heterooligomeric complexes with other CaMK2 subunits (CaMK2β, CaMK2γ and CaMK2δ)^23^. To profile the translational landscape of *Camk2a*⁺ cells, we delivered a cocktail of viral vectors encoding cre recombinase under short *Camk2a* promoter (0.4 kb)^24^ and cre-dependent fluorescent ribosome tag^25^ into the PL. Following three weeks of viral-mediated gene expression (**Fig. 2a**), mice underwent PTC, and PL tissue was collected at 1 hour or 4 hours post-training for translating ribosome affinity purification (TRAP)^26^, followed by bulk RNA sequencing to capture ribosome-associated mRNAs specifically from *Camk2a*⁺ cells (**Fig. 2b**). Box-Only mice, which were exposed to the training context without CS–US pairings, served as controls. Behaviorally, Trained mice exhibited a significant increase in freezing during the second CS–US pairing, whereas Box-Only controls maintained minimal baseline freezing during dummy CS epochs (**Fig. 2c**).

**Figure 2.**
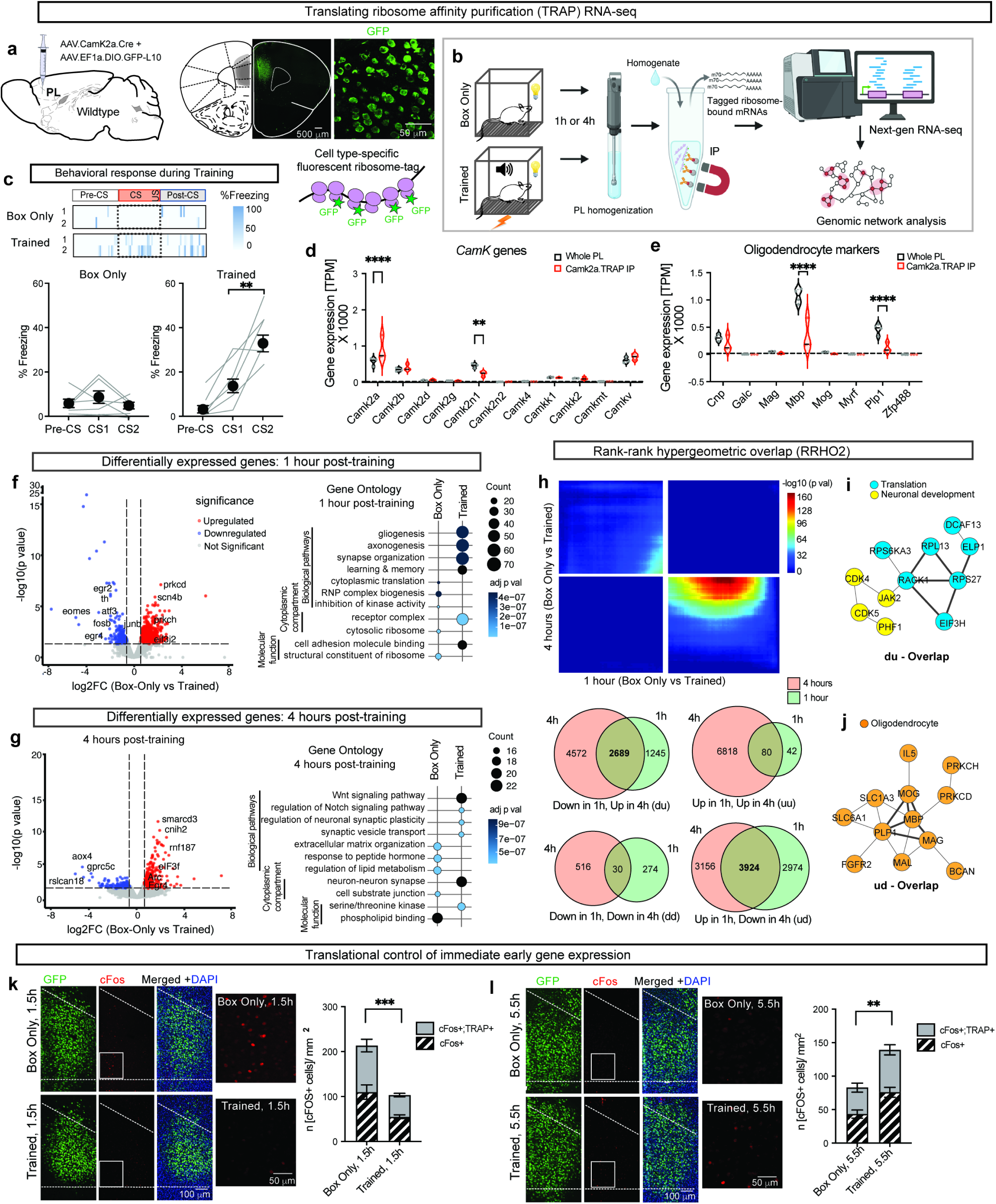
Biphasic translational reprogramming during memory consolidation. **a)** Viral delivery of a fluorescent ribosome tag to Camk2a⁺ cells in the prelimbic (PL) cortex. Scale bar = 500 μm, 50 μm. **b)** PL tissue from Box-Only and PTC-trained mice was collected at 1 h or 4 h post-training for translating ribosome affinity purification followed by RNA sequencing (TRAP-seq). **c)** Representative heatmap showing freezing response during pre-CS, CS, and post-CS epochs (30s each) in Box Only and PTC trained mice in Context A. Trained mice exhibited significantly increased freezing during the second CS–US pairing, whereas Box-Only controls remained at baseline. Box Only: F (1.777, 10.66) = 1.204; Trained: F (1.392, 8.353) = 33.83; n=7 mice/group. **d)** Transcripts per million (TPM) comparison revealed increased *Camk2a* and reduced *Camk2n1* expression in trained samples relative to the global PL transcriptome. Marker gene (*Camk*) X TRAP: F (10, 55) = 4.770; n=3-4 mice/ group. **e)** Oligodendrocyte markers were detected but significantly depleted in Camk2a⁺ TRAP samples compared to whole PL. Marker gene (Oligodendrocytes) X TRAP: F (7, 40) = 14.59; n=3-4 mice/ group. **f)** Volcano plot displaying the total DEGs in the trained TRAP samples compared to Box-Only control group at 1h timepoint (p < 0.05, |log₂FC| > 0.58). At 1 h post-training, DEGs showed repression of neuronal and synaptic transcripts with concurrent induction of oligodendrocyte-associated genes; GO analysis indicated enrichment for gliogenesis and axonogenesis, alongside suppression of translation-related pathways. n=3 animals/group. **g)** Volcano plot exhibited the total DEGs in the trained TRAP samples compared to Box-Only control group at 4h timepoint (p < 0.05, |log₂FC| > 0.5). At 4 h, translational programs shifted toward induction of synaptic plasticity and Wnt signaling, while Box-Only controls were enriched for lipid metabolism and extracellular matrix organization. n=3 mice/group. **h–j)** Rank-rank hypergeometric overlap (RRHO2) analysis uncovered reciprocal gene programs across time points, with early upregulation of glial-associated genes and delayed induction of translation and neurodevelopmental modules. **k–l)** Immunohistochemistry revealed decreased cFos⁺TRAP⁺ cells at 1.5 h and increased neuronal activation at 5.5 h following training. n=3 mice/group. **Statistical tests: c)** One-way ANOVA with Bonferroni post-hoc test; **d, e)** Two-way ANOVA with Bonferroni post-hoc test; **k, l)** Unpaired t-test. **p<0.01, ****p<0.0001.

Quality control analyses confirmed the robustness of the RNA-seq dataset. Biological replicates exhibited strong within-group correlation, clustered consistently across samples, and segregated appropriately by principal component analysis, with PC1 and PC3 explaining 44.0% and 13.7% of the variance at 1 hour, and PC1 and PC2 explaining 66.6% and 16.8% at the later time point (**Extended Data Fig. 4a-c**). MA plots further demonstrated that majority of transcripts clustered tightly around the midline, with only a small subset showing significant differential expression (**Extended Data Fig. 4d**). Differential TPM analysis revealed a marked upregulation of *Camk2a* transcript together with a concomitant reduction of its endogenous inhibitor *Camk2n1* in Trained TRAP samples compared to the bulk PL transcriptome (**Fig. 2d**). Other *Camk2* subunits were not differentially expressed in TRAP samples, however *Camk2b* and *Camkv* transcripts were expressed above threshold. Canonical neuronal markers such as *Slc17a7, App, Map2, Nrn1, Sv2a, Nrgn,* and *Snap25* were consistently expressed above threshold in *Camk2a*⁺ TRAP samples (**Extended Data Fig. 5a**). Given the established role of CaMK2β in oligodendrocyte maturation and the prior observation that *Camk2a* is also expressed at low levels in oligodendrocytes^27^, we next examined the expression of oligodendrocyte markers in our dataset. Several myelination-associated transcripts, including *Cnp, Mbp,* and *Plp1*, were robustly detected above expression threshold, whereas canonical astrocyte markers were not (**Extended Data Fig. 5b**). Although their expression levels were lower than in the whole PL transcriptome, their presence in the *Camk2a*⁺ TRAP dataset indicates that the captured population includes both neurons and oligodendrocytes (**Fig. 2e; Extended Data Fig. 5c**).

Differential expression analysis at the 1h time point identified 822 transcripts as significantly upregulated and 423 as downregulated in Trained TRAP samples relative to Box Only controls (p < 0.05, |log₂FC| > 0.58), with volcano plots revealing marked repression of immediate early genes (*Fosb, Egr2, Junb*) and synaptic plasticity–associated transcripts (*Calb2, Th, Atf3*), alongside robust induction of oligodendrocyte-enriched genes (*Mag, Mbp, Mal*) and translation factor eIF3 subunit *eif3j*, a gene implicated in regulating protein synthesis of select mRNAs^28,29^ (**Fig. 2f**, left; **Supplementary Table 1**). Consistent with this, Gene Ontology (GO) analysis showed enrichment of gliogenesis, axonogenesis, and synapse organization among upregulated genes and cytoplasmic translation and ribonucleoprotein complex biogenesis among downregulated genes, indicating rapid suppression of neuronal translational programs in the prelimbic cortex (PL) during early consolidation with concurrent activation of oligodendrocyte-associated pathways (**Fig. 2f**, right; **Supplementary Table 2**). In addition, a discrete but coherent signature of PERK-eIF2α signaling that mediates both integrated stress response^30^ and unfolded protein response (UPR)^31^ was detectable within the dataset. Canonical markers of endoplasmic stress and UPR activation - including *Hspa5* (BiP), *Pdia4*, *Pdia3*, *Hyou1*, and *Os9* - were robustly represented (**Extended Data Fig. 5d**), indicating engagement of ER proteostasis pathways downstream of PERK activation. Additional components associated with translational stress adaptation and proteostatic remodeling, such as *Emc1*, *Stt3b*, and *Clptm1*, further support the presence of a coordinated stress-response program.

By contrast, at the 4h time point, 304 genes were upregulated and 116 downregulated, with strong induction of *Arc* and *Egr4* and enrichment of Wnt and Notch signaling, synaptic plasticity, and synaptic vesicle transport pathways, while Box Only control-enriched transcripts reflected extracellular matrix organization, peptide hormone responses, and lipid metabolism (**Fig. 2g; Supplementary Table 3, 4**). Rank–rank hypergeometric overlap (RRHO2) analysis^32^ confirmed a reciprocal translational program across timepoints (**Fig. 2h; Supplementary Table 5**), and network analysis in Cytoscape identified a translation-centered module among genes downregulated at 1 h and upregulated at 4 h (e.g., RPS6KA3, RPL3, RPS27, RACK1, EIF3H, ELP1, DCAF13), alongside neurodevelopmental regulators (CDK4, CDK5, JAK2, PHF1) (**Fig. 2i**), whereas the inverse gene set formed a myelination-centric network enriched for oligodendrocyte plasticity–associated genes (MBP, MOG, MAG, PLP1, SLC1A3, SLC6A1, FGFR2, IL5, PRKCH, PRKCD, BCAN) (**Fig. 2j**), revealing a biphasic reorganization of the PL translatome characterized by early translational repression and glial activation followed by delayed neuronal translation, a pattern corroborated by immunohistochemistry showing reduced cFos protein expression at 1.5 h and increased expression at 5.5 h post-training (**Fig. 2k, l**).

### Cap-dependent translation governs memory reconsolidation and systems stabilization

It is well established that recent memory consolidation requires *de novo* protein synthesis via both cap-dependent and cap-independent mechanisms in subcortical limbic regions such as the amygdala^4,33^ and hippocampus^34–36^. In light of our discovery of a biphasic translational program in the prelimbic cortex (PL), we next sought to test whether 7-methylguanosine (m7G)^37^ cap-dependent translation in *Camk2a*-expressing cells plays a causal role in remote memory consolidation, both in the presence and absence of recent retrieval. To directly address this question, we implemented a loss-of-function strategy that suppressed cap-dependent translation in PL *Camk2a*⁺ cells not only during early consolidation but also throughout the systems consolidation window spanning several weeks. To achieve sustained and cell type–specific translational inhibition, we employed an intersectional chemogenetic approach using a knock-in mouse line that enables Tet- and Cre-dependent expression of an *eif4e*-targeting short hairpin RNA embedded within a miR-30 backbone^4,38^. This design allowed stable depletion of eIF4E (4Ekd) selectively in PL *Camk2a*⁺ cells under doxycycline-off (Off-Dox) conditions initiated 10 days before training (**Fig. 3a**, top & right; **Fig. 3b**). Because eIF4E is a rate-limiting determinant of cap-dependent translation initiation^2^, this manipulation provides a persistent and targeted reduction of protein synthesis across both early and late phases of memory consolidation. The 4Ekd strategy had no effect on spontaneous locomotion or thigmotaxis in the open field test, suggesting that eIF4E levels in PL *Camk2a*⁺ cells do not modulate baseline anxiety (**Extended Data Fig. 6a-c**).

**Figure 3.**
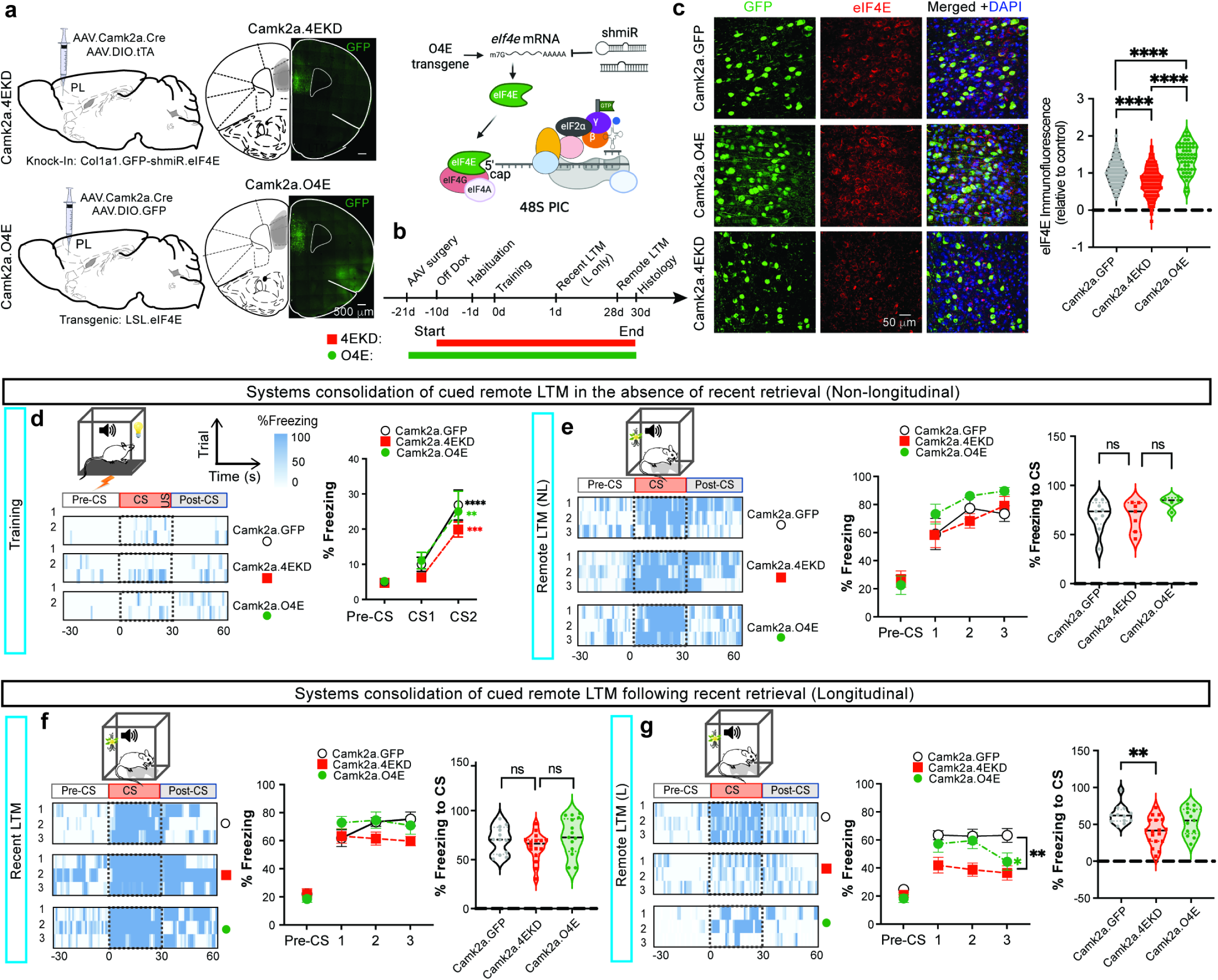
Cap-dependent translation governs memory reconsolidation and systems stabilization. **a**) Schematic illustrating the (chemo)genetic strategies to deplete or overexpress eIF4E, i.e., 4Ekd or O4E, to manipulate levels of 48S preinitiation complex (PIC). Knock-in mice, Col1a2.GFP-shmiR.eIF4E, were injected with AAV9.Camk2a.cre and AAV9.Eef1a.DIO.tTA into the PL to knockdown eIF4E (4EKD) in PL *Camk2a*⁺ cells in a Cre-and-tet dependent manner, whereas transgenic LSL-eIF4E mice were injected with AAV9.Camk2a.cre. **b)** Timeline for eIF4E manipulation and PTC longitudinal and non-longitudinal remote LTM testing. **c)** Immunostaining of eIF4E protein in PL *Camk2a*⁺ cells in eIF4E knockdown (Camk2a.4Ekd) and eIF4E overexpression (Camk2a.O4E) groups compared to control (Camk2a.GFP) group mice. F (2, 720) = 94.21; n = 55-342 cells from 3 mice/group. **d)** PTC Training for Camk2a.GFP, Camk2a.4Ekd and Camk2a.O4E groups show equivalent learning of the CS-US associations, both in representative heatmaps (showing freezing response during pre-CS, CS, and post-CS epochs) and X-Y plot. Time x Genotype: F (2, 42) = 0.2519; n=13-17 mice/group. **e)** CS-evoked freezing response during single timepoint NL-remote LTM was not affected in Camk2a.GFP, Camk2a.4Ekd and Camk2a.O4E groups – as evident in the representative heatmaps, X-Y plot showing freezing response for each CS (CS x Genotype: F (4, 36) = 0.5577) and violin plot showing average freezing response for all CS’s (F (2, 18) = 1.913). n=5-9 mice/group. **f)** CS-evoked freezing response during recent LTM is intact in all three groups (Camk2a.GFP, Camk2a.4Ekd and Camk2a.O4E), as evident in the representative heatmaps, X-Y plot showing freezing response for each CS (CS X Genotype: F (4, 74) = 2.375) and violin plot showing average freezing response for all CS’s (F (2, 38) = 1.499). n=12-15 mice/group. **g)** When longitudinally tested, the L-remote LTM was significantly impaired for both Camk2a.4Ekd and Camk2a.O4E groups compared to control Camk2a.GFP mice. Representative heatmaps showing freezing response during pre-CS, CS, and post-CS epochs (30s each), and X-Y plot showing freezing response for each CS (Genotype: F (2, 38) = 7.398) and violin plot showing average freezing response for all CS’s (F (2, 40) = 6.614) show the effect of eIF4E manipulation on reconsolidation. n=12-17 mice/group. **Statistical tests: c, e** (right)**, f** (right)**, g** (right)) One-way ANOVA with Bonferroni post-hoc test; **d** (right)**, e** (middle)**, f** (middle)**, g** (middle)) Two-way RM ANOVA with Bonferroni post-hoc test. *p<0.05, **p<0.01, ***p<0.001, ****p<0.0001, ns not significant.

In a complementary approach, we utilized the transgenic *Rosa26-eif4e* mouse strain^39^ to Cre-conditionally overexpress eIF4E (o4E) in PL *Camk2a*⁺ cells (**Fig. 3a**, bottom & right) in adult life following surgery (**Fig. 3b**). Both 4Ekd and o4E strategies were effective at bidirectionally modulating the abundance of eIF4E protein in PL *Camk2a*⁺ cells (**Fig. 3c**). Behaviorally, neither eIF4E knockdown nor overexpression altered acquisition during training (**Fig. 3d**) or affected single time-point NL-remote LTM (**Fig. 3e**). However, in the longitudinal framework - although recent LTM was preserved across groups (**Fig. 3f**), memory maintenance following retrieval-induced destabilization, i.e., reconsolidation^40,41^, was significantly more vulnerable under both eIF4E depletion and overexpression (**Fig. 3g**). To determine whether eIF4E depletion impairs reconsolidation in intermediate timepoint as well, animals were switched to Off-Dox diet beginning 7 days after training, underwent retrieval at 14 days, and were tested for longitudinal remote LTM at 28 days. While LTM at 14 days was unaffected, remote LTM was markedly impaired in the 4Ekd group relative to GFP controls (**Extended Data Fig. 7a-c**). Together, these results demonstrate that precise regulation of cap-dependent translation following retrieval across memory age is critical for reconsolidation and systems-level stabilization of memory traces.

### Chemogenetic activation of PKR reprograms translation in PL *Camk2a*⁺ cells

Given that cap-dependent translation in PL Camk2a⁺ cells does not causally contribute to memory consolidation, we next examined the molecular and behavioral consequences of manipulating cap-independent translation. To this end, we employed chemogenetic iPKR, a knock-in system that enables rapid and reversible, cell type-specific inhibition of protein synthesis via drug-inducible phosphorylation of eIF2α (Ser51)^4^. iPKR mice were injected with a viral cocktail to co-express iPKR and the fluorescent ribosome tag in PL *Camk2a*⁺ cells (iPKRTRAP) and implanted with a lateral ventricle cannula for the iPKR-inducer drug (ASV) infusion immediately after training. To assess the change in translation landscape following iPKR activation, cell type-specific translating ribosome affinity purification (TRAP) was performed 1 h later, followed by bulk RNA-seq of ribosome-associated mRNAs (**Fig. 4a**). Control (Box Only) iPKRTRAP also received ASV infusion immediately after context exposure. Trained and Box Only iPKRTRAP mice were then compared with the corresponding TRAP samples lacking iPKR. RNA-seq quality assessments confirmed high dataset integrity: biological replicates showed strong within-group concordance, samples clustered appropriately, and principal component analysis showed that PC1 and PC2 accounted for 60.2% and 17.1% of the variance for Box Only mice and 71.1% and 15.7% for Trained mice (**Extended Data Fig. 8a-c**). MA plots demonstrated that most transcripts were centered near log₂ fold-change zero, with a limited subset showing significant deviations consistent with genuine differential expression (**Extended Data Fig. 8d**). Notably, TPM analysis revealed elevated expression of *Atf4* - the canonical mediator of the integrated stress response^30^ - in trained iPKRTRAP samples relative to TRAP controls (**Fig. 4b**) along with downregulation of the plasticity-related genes - *Camk2a* (**Fig. 4c**) as well as *Arc, Atp1a3, Calb1* and *Egr1* (**Extended Data Fig. 9**).

**Figure 4.**
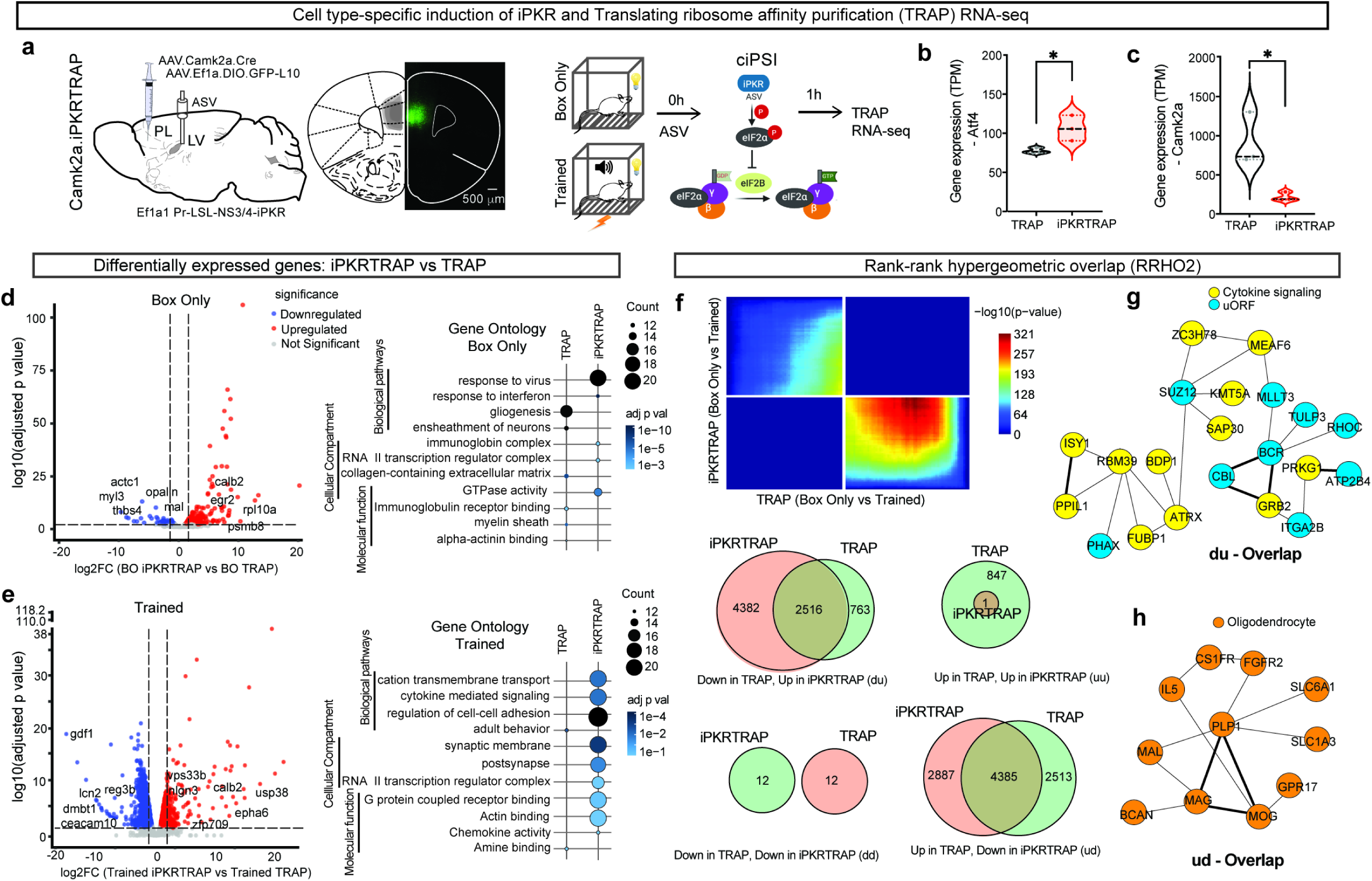
Chemogenetic activation of PKR reprograms translation in PL *Camk2a*⁺ cells. **a)** Schematic illustration of viral cocktail (AAV.Camk2a.cre and AAV.EF1a.DIO.GFP-L10) into PL of iPKR mice to co-express iPKR and a fluorescent ribosome tag in PL *Camk2a*⁺ cells (iPKRTRAP). The Box Only and PTC-trained iPKRTRAP mice received Asunaprevir (ASV) infusion via lateral ventricle cannula immediately after PTC training. PL tissue was harvested 1h later for Translating ribosome affinity purification (TRAP) followed by RNA-sequencing (TRAP-Seq). Scale bar = 500 μm. **b)** Transcripts per million (TPM) analysis revealed a robust upregulation of *Atf4* transcript in trained iPKRTRAP samples relative to the trained TRAP group. n=3 mice/ group. **c)** left: Volcano plot exhibiting the total DEGs identified in the iPKRTRAP samples compared to TRAP Box Only controls (adj. p < 0.05, |log₂FC| > 0.58). right: Gene ontology (GO) enrichment analysis identified upregulated transcripts involved in categories related to viral response and interferon signaling in iPKRTRAP Box Only samples compared to TRAP Box Only controls (FDR adjusted p < 0.05). **d)** Volcano plot exhibiting the total DEGs identified in the iPKRTRAP Trained samples compared to TRAP Trained controls (adj. p < 0.05, |log₂FC| > 0.58). **e)** GO enrichment analysis identified upregulated transcripts involved in cation transmembrane transport and cytokine-mediated signaling in iPKRTRAP Trained samples compared to TRAP Trained controls (FDR adjusted p < 0.05). **f)** Rank-Rank Hypergeometric Overlap (RRHO2) method was used for gene set enrichment analysis using lists of identified DEGs in TRAP and iPKRTRAP samples (Box vs. PTC-trained) to identify statistically significant transcriptional concordant and discordant overlaps between the two groups. Venn diagram showing the total number of identified upregulated, downregulated and overlapping genes in TRAP vs. iPKRTRAP groups under Box Only vs PTC-trained conditions. **g)** Gene network analysis of discordant genes (downregulated in TRAP and upregulated in iPKRTRAP, 1 hour) revealed gene module related to cytokine signaling and transcripts bearing upstream open reading frames. **h)** Gene network analysis of discordant genes (upregulated in TRAP and downregulated in iPKRTRAP, 1 hour) revealed gene module related to oligodendrocyte markers. n=3 mice/group. **Statistical test:** Unpaired t-test. *p<0.05; **c, d, e, f)** statistical analyses outlined in Methods section.

Differential gene expression analysis showed that in the Box Only condition, 267 transcripts were upregulated, and 164 were downregulated in iPKRTRAP relative to TRAP controls (**Fig. 4d**, left), whereas in the Trained condition, 724 transcripts were upregulated and 1276 were downregulated (adj. p <0.05, |log₂FC| > 0.58) (**Fig. 4e**, left; **Supplementary Table 6, 7**). Gene ontology (GO) enrichment analysis of upregulated transcripts in both conditions showed strong enrichment for interferon-related signaling pathways, consistent with the known role of endogenous PKR in activating antiviral responses upon detection of viral RNA^42^. In the Box Only comparison, this included genes such as *Bst2, Dtx3l, Eef2ak2, Gbp2*, and *Ifit1* (**Fig. 4d**, right), whereas the Trained comparison highlighted *Ackr1, Adipor2, Apoa1, C1qtnf*, and *Ccl12* (**Fig. 4e**, right). By contrast, downregulated transcripts in iPKRTRAP were enriched for GO terms linked to myelination (e.g., *Adora2a, Aspa, Csf1r* and *Cspg4*) for Box Only comparisons and adult behavior (e.g., *Atg7, Nlgn3, Chrna7*, and *Htr2a)* for Trained comparisons. In the Trained condition, additional enrichment was observed for regulation of cation transmembrane transport (e.g., *Adrb1, Atp2b4, Cacna1a, Calca, Drd2, Grin2d, Nlgn2* and *Slc30a1*) and cell adhesion (e.g., *Actb, Apoa1, Ccdc88b, Ccl19* and *Ccl2*), suggesting that iPKR activation disrupts both immune-related and synaptic signaling programs during consolidation (**Supplementary Table 8, 9**). RRHO2 analysis for TRAP (Box Only vs Trained) and iPKRTRAP (Box Only vs Trained) samples (**Fig. 4f**;) further underscored the discordance between conditions (**Supplementary Table 10**). Notably, iPKRTRAP Trained samples showed selective translational upregulation of cytokine-related genes and uORF-containing transcripts, including CBL, BCR, MLLT3, and TULP3 (**Fig. 4g**), consistent with iPKR-dependent, gene-selective translation during ISR activation. Conversely, a gliogenesis-associated gene network was upregulated in TRAP Trained samples but suppressed in iPKRTRAP Trained samples (**Fig. 4h**), indicating that iPKR activation antagonizes training-induced glial remodeling at the translational level 1h post-training.

### Early cap-independent translation gates recent–remote memory tradeoff

We next assessed the behavioral impact of acute iPKR activation in PL Camk2a⁺ cells during the early consolidation window (0-3 h post-training) on long-term memory formation (**Fig. 5a**). Control Camk2a.GFP mice received identical post-training ASV infusions. Both Camk2a.GFP and Camk2a.iPKR mice showed comparable acquisition during training, as evidenced by a significant increase in freezing between the first and second CS (**Fig. 5b**), indicating intact learning. When tested for recent long-term memory, Camk2a.iPKR mice exhibited a pronounced impairment relative to GFP controls (**Fig. 5c**). This deficit - absent when cap-dependent translation was suppressed (**Fig. 3f**) - reveals a critical requirement for rapid, cap-independent translation of myelination- and unfolded protein response–related genes in PL *Camk2a*⁺ cells during early consolidation. Strikingly, the same animals displayed a significant enhancement in memory strength at the remote timepoint (L-Remote LTM) (**Fig. 5d**). Together, these findings reveal an inverse relationship between recent and remote memory expression, suggesting that disruption of recent memory retrieval in iPKR mice occludes retrieval-induced forgetting^43^ and ultimately promotes stronger memory expression at remote timepoints. In contrast, when non-longitudinal (NL) remote memory was tested at a single timepoint (28 days post-training), Camk2a.iPKR mice showed no difference in memory strength relative to controls (**Extended Data Fig. 10a, Fig. 5e**), indicating that early cap-independent translation in PL *Camk2a*⁺ cells is not required for remote memory formation per se when assessed without prior retrieval. However, sustained enhancement of protein synthesis achieved through homozygous mutation of the iPKR target site on eIF2α (S51A)^4,44^ (**Fig. 5f**), which prevents phosphorylation-mediated translational repression, resulted in significantly elevated freezing at the remote test (**Extended Data Fig. 10b, c, Fig. 5g**). Together, these findings identify eIF2α-dependent translational control as a determinant of delayed - rather than early - systems-level consolidation of remote cued threat memories.

**Figure 5.**
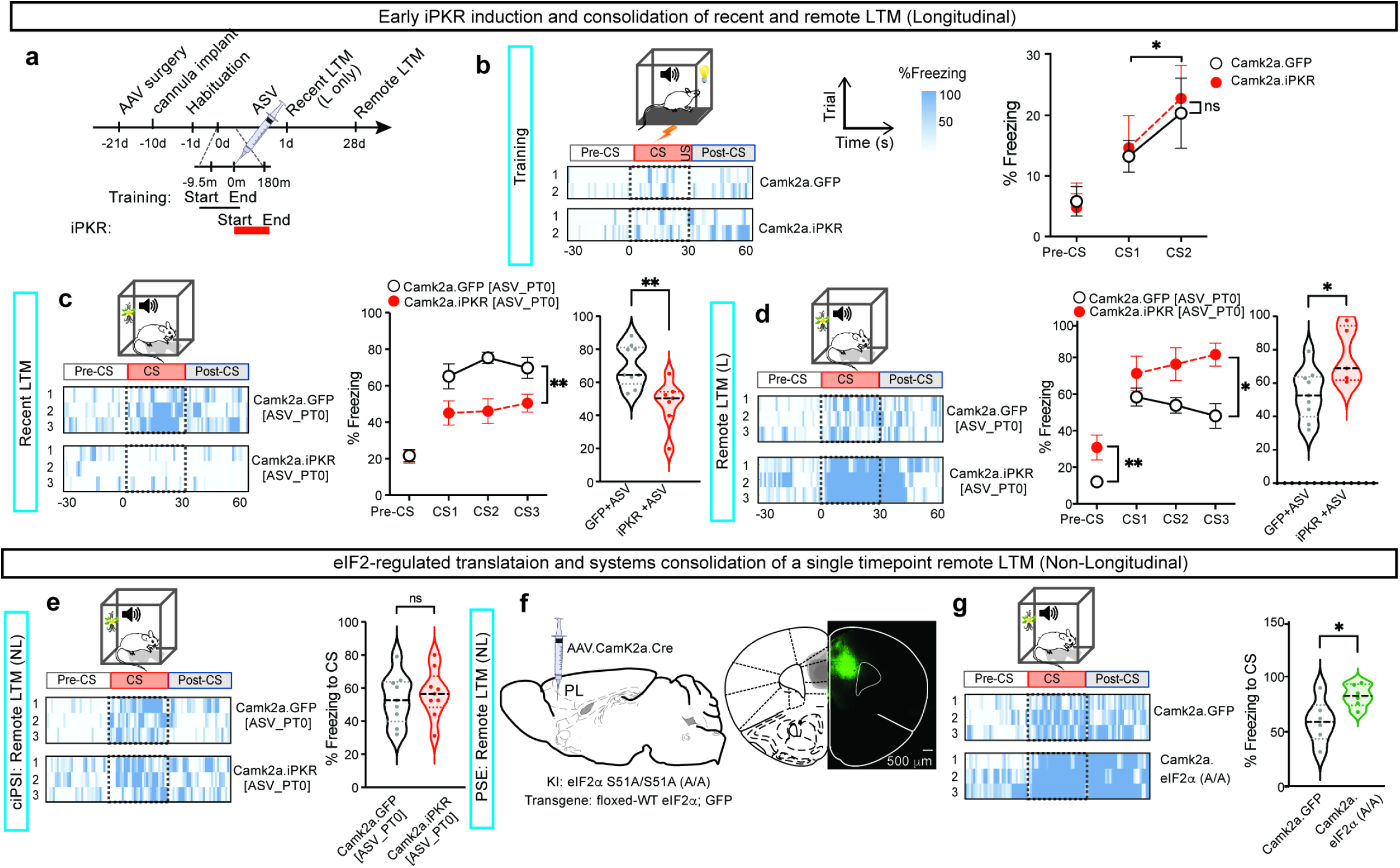
Early cap-independent translation gates recent–remote memory tradeoff. **a)** Schematic illustration of the experimental timeline for Camk2a.iPKR and Camk2a.GFP control mice – ASV infusion is carried out immediately after training for both groups, and remote memory retention is tested in either longitudinal or non-longitudinal framework. **b)** left: Representative heatmaps showing freezing response during pre-CS, CS, and post-CS epochs (30s each) in Camk2a.GFP and Camk2a.iPKR mice. right: X-Y plot showing similar learning capacity during PTC training for both groups (Time X Genotype: F (1, 14) = 0.02096), as revealed by robust increase in % freezing to the second CS (F (1, 14) = 4.746). n=8 mice/group. **c)** left: Representative heatmaps showing freezing response during pre-CS, CS, and post-CS epochs (30s each) at recent LTM, tested 1 day after training. middle: X-Y plot showing a significant decrease in CS-evoked freezing response at recent LTM in Camk2a.iPKR mice compared to Camk2a.GFP controls, while pre-CS freezing response is unchanged (Genotype: F (1, 14) = 11.47). right: Violin plot showing mean CS-evoked freezing response in Camk2a.iPKR and Camk2a.GFP mice. n=5-9 mice/group. **d)** left: Representative heatmaps showing freezing response during pre-CS, CS, and post-CS epochs at longitudinally tested remote LTM (L-remote LTM). middle: X-Y plot showing a significant increase in L-remote LTM in Camk2a.iPKR mice compared to Camk2a.GFP controls (Time X Genotype: F (2, 24) = 4.593). Freezing response during pre-CS is also increased in the Camk2a.iPKR mice. right: Violin plot showing mean CS-evoked freezing response in Camk2a.iPKR and Camk2a.GFP mice. n=5-9 mice/group. **e)** left: Representative heatmaps showing freezing response during pre-CS, CS, and post-CS epochs at single-timepoint remote LTM (NL-remote LTM) in Camk2a.iPKR and Camk2a.GFP mice. right: Violin plot showing mean CS-evoked freezing response during NL-remote LTM in Camk2a.iPKR and Camk2a.GFP groups. **f)** Schematic illustration showing injection of AAV9.Camk2a.Cre into the PL of knock-in/transgenic homozygous eIF2α phosphomutant (S51A) mice. Immunostained prefrontal brain section showing mutant eIF2α (A/A) and GFP reporter expression in the PL. Scale bar = 500 μm. **g)** Representative heatmaps showing freezing response during pre-CS, CS, and post-CS epochs at single-timepoint remote LTM (NL-remote LTM) for Camk2a.eIF2α (A/A) and Camk2a.GFP mice. right: Violin plot showing mean CS-evoked freezing response during NL-remote LTM for Camk2a.eIF2α (A/A) and Camk2a.GFP groups. **Statistical tests: b, c** middle, **d** middle) RM Two-way ANOVA with Bonferroni post-hoc test; **c** right, **d** right, **e** right, **f** right) Unpaired t-test. *p<0.05, **p<0.01, ns not significant.

## Discussion

Our findings support a model in which the temporal organization of translational control within the prelimbic cortex plays a central role in shaping the durability of threat memories (**Figure 6a, b**). Under a non-longitudinal design in which animals were probed only once at the 28-day timepoint, learning engaged a biphasic translational program: an early phase dominated by ER-stress–mediated suppression of protein synthesis coupled with pronounced oligodendrocyte plasticity, and a later phase marked by synaptic growth and stabilization. These sequential molecular states appear to work in concert to promote the long-term persistence of remote memory. Notably, both recent and remote consolidation depend on eIF2α-regulated, cap-independent translation, underscoring the importance of this pathway for systems-level stabilization of learned associations. In contrast, when a memory was retrieved during the recent period, the act of reactivation induced destabilization of the underlying trace, triggering a reconsolidation process that operated through a distinct, eIF4E-dependent translational mechanism. This pathway only partially restored memory stability, revealing a separation between consolidation- and reconsolidation-linked modes of protein synthesis. Importantly, even though remote memories did not exhibit passive decay over the same interval, retrieval itself introduced a vulnerability: the destabilization–reconsolidation cycle weakened memory persistence, highlighting retrieval-induced forgetting as a key determinant of long-term memory strength.

**Figure 6.**
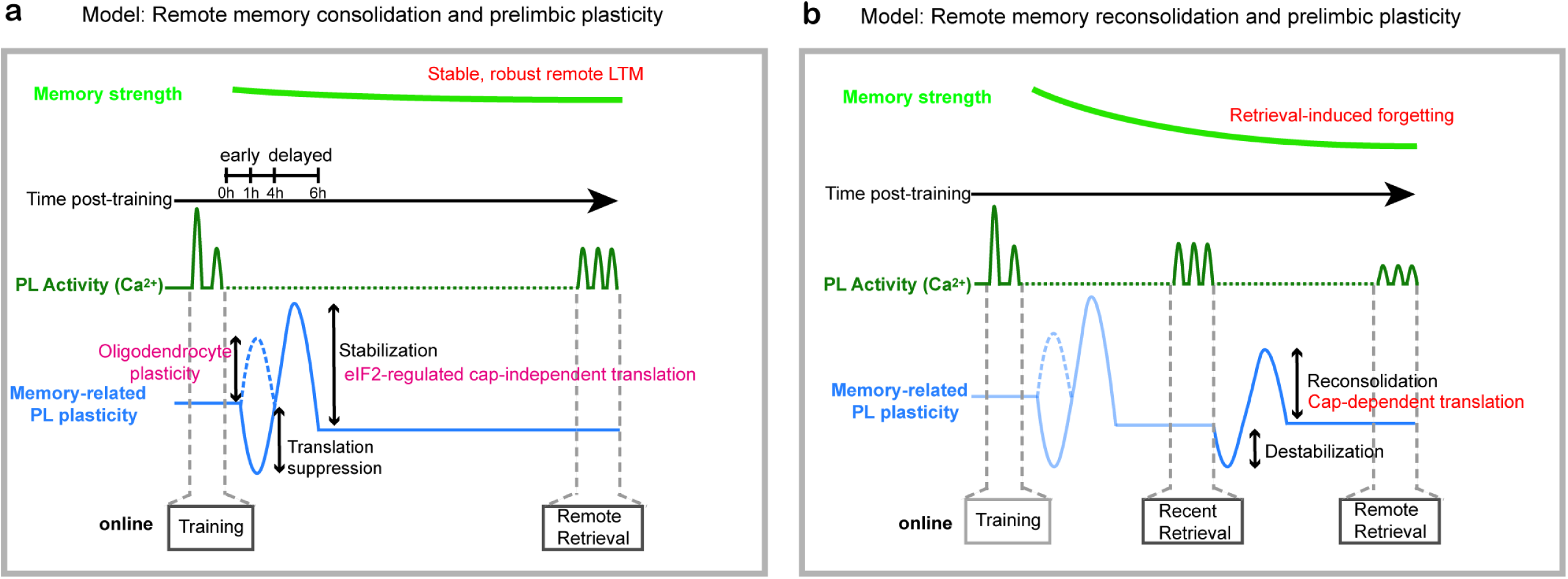
Schematic representations illustrate proposed mechanisms linking remote memory strength and memory-related plasticity in mice after post-training interventions. **a**) Model illustrating the evolution of memory strength and prelimbic cortical plasticity under the non-longitudinal testing design, in which mice were probed only once for remote memory at 28 days after training. Learning initiates a biphasic translational program: an early wave marked by robust ER-stress–driven suppression of protein synthesis together with heightened oligodendrocyte plasticity, followed by a later phase characterized by synaptic growth and stabilization. Together, these temporally structured processes support long-term memory persistence. Both recent and remote consolidation require eIF2-dependent, cap-independent translation. **b)** In contrast, recent retrieval reactivates and destabilizes the memory trace, initiating a reconsolidation process that relies on an eIF4E-dependent translation program to partially—but not fully—restore memory stability. Although remote memories do not undergo passive time-dependent decay within this window, retrieval-induced weakening mechanisms nevertheless erode their long-term persistence.

Although our viral approach targeted all *Camk2a*-expressing cells within the prelimbic cortex, thereby capturing both neurons and oligodendrocytes, this broader inclusion proved unexpectedly informative. The early consolidation phase, marked by global suppression of translation, coincided with pronounced oligodendrocyte plasticity and increased myelination - an adaptation that would be expected to enhance conduction velocity and optimize information transfer along pyramidal neuron axons^45,46^. This temporal arrangement is striking; while translation in pyramidal neurons is largely paused, the circuit simultaneously upgrades its structural “infrastructure,” and only after this myelination-driven increase in throughput does translation resume to support synaptic growth programs, including those engaging Wnt- and Notch-dependent pathways^47,48^. The delayed translational wave may also reflect systems-level coordination between the PL cortex and subcortical mnemonic structures, consistent with classical evidence for temporally delayed consolidation processes^49,50^. Memory reactivation during quiet wakefulness is known to involve coupling between cortical spindles and hippocampal or amygdala sharp-wave ripples^51–53^. Notably, Steadman et al (2020)^54^ demonstrated that blocking learning-induced oligodendrogenesis - mirroring the phenomenon observed here - disrupts the learning-associated increase in hippocampal ripple–cortical spindle coupling. Such replay events preserve the spatiotemporal structure of prior experience and are thought to promote systems consolidation. The convergence of this replay window with the shift from translational suppression to activation suggests that reinstatement of memory traces may act as a trigger - or at minimum a permissive context - for the subsequent wave of protein synthesis that stabilizes long-term memory. An important future direction will be to determine whether remote persistence depends on multiple delayed, offline translational waves during non-REM sleep or quiet wakefulness across the days and weeks following learning.

Finally, the retrieval-induced forgetting observed here was robust under a moderate Pavlovian threat-conditioning paradigm (two tone–shock pairings, 0.4 mA footshock). Whether stronger or more traumatic learning conditions buffer against this vulnerability - and whether such buffering is accompanied by altered translational dynamics - remains an open question. Addressing these issues will be essential for understanding how retrieval history, translational regulation, and systems-level plasticity interact to shape the longevity of emotional memories.

## Author contributions

Conceptualization: P.S. and J.I. Methodology: J.I., S.K., S.L., A.S., L.P., H.S., A.G. and A.W. Investigation: J.I., S.K., S.L., A.S., L.P., H.S., A.G. and A.W. Funding acquisition: P.S. Supervision: P.S. Writing – original draft: P.S. and J.I. Writing – review & editing: J.I., S.K., S.L., A.S., L.P., H.S., A.G., A.W., B.X. and P.S.

## Acknowledgements

We thank Alex Fraser, Ananya Chaubal, Keith Yeung, and Stephanie Chu for technical assistance; Dr. David Carlson (Institute for Advanced Computational Science, Stony Brook University), Dr. Nicolas Robine (NY Genome Center), Dr. Jeffrey Dupree (Virginia Commonwealth University), and all Shrestha Lab members for critical feedback and discussions. AAV.Camk2a.Cre (pENN.AAV.CamKII0.4.Cre.SV40) was a gift from James M. Wilson (Addgene viral prep #105558-AAV9; http://n2t.net/addgene:105558; RRID:Addgene_105558). AAV.CAG.GCaMP6f (pAAV.CAG.GCaMP6f.WPRE.SV40) was a gift from Douglas Kim & GENIE Project (Addgene viral prep #100836-AAV1; http://n2t.net/addgene:100836; RRID:Addgene_100836). AAV.EF1a.DIO.GFP-L10 (pAAV-FLEX-EGFPL10a) was a gift from Nathaniel Heintz & Alexander Nectow & Eric Schmidt (Addgene viral prep #98747-AAV1; http://n2t.net/addgene:98747; RRID:Addgene_98747). P.S. is supported by National Institute of Mental Health grant R01 MH132795, Alfred P Sloan Foundation fellowship FG-2022-94485. S.L. is supported by Scholars in Biomedical Sciences Program (T32GM148331). B.X. is supported by National Institutes of Mental Health grant R01 MH125187.

## Materials and Methods

### Animals

Both male and female mice aged 3-6 months old were used in all experiments. All procedures involving the use of animals were performed in accordance with the guidelines of the National Institutes of Health and were approved by the University Animal Welfare Committee of Stony Brook University. Animals were group-housed in a 7.5 in x11.5 in cage with corn cob bedding and enrichment (shredded paper) and were provided with food and water *ad libitum*. Animals were maintained in a 12-hour (light on 7 AM - light off 7 PM) light/dark cycle in the institutional animal care facility at stable temperature (78°F) and humidity (50%). Wildtype C57Bl/6J mice were purchased from Jackson Laboratory (stock #000664). Chemogenetic iPKR strain^1^ was developed by Shrestha & Ayata, and received from Dr Nathaniel Heintz (The Rockefeller University); Col1a1TRE GFP.shmir4E.389^2^ mice were a gift from Dr Jerry Pelletier (McGill University) and transgenic homozygous Eif2s1 (S51A); CAG Prfl.Eif2s1.fl.GFP mice^3^ (that is, eIF2α (A/A)) were a gift from Dr Randal Kaufman (Sanford Burnham Prebys Medical Discovery Institute). Rosa26-eIF4E mice^4^ were generated in Baoji Xu’s lab (The Scripps Research Institute Florida). All mice were backcrossed with the C57Bl/6J strain for at least five generations.

### Drugs and chemicals

ASV (MedChemExpress, Catalog #HY-14434) was dissolved in dimethylsulfoxide (DMSO) for a stock concentration of 10 mM and diluted in sterile saline to 100 nM. Two microliters of this drug were intracranially infused into the left lateral ventricle (−0.50 mm anteroposterior (AP), −1.10 mm mediolateral (ML) and −2.20 mm dorsoventral (DV)). Doxycycline was prepared in rodent chow (Bio-Serv, Catalog #F4159) at 40 mg/kg, and this diet was provided to the mice in 4Ekd group for 2 or 4 weeks following surgery. Transcardial perfusion was performed using 4% paraformaldehyde (PFA). A stock solution of aqueous 32% paraformaldehyde PFA (EMS, #15714) was freshly diluted to 4% PFA in 0.1 M PBS for transcardial perfusion and post-fixation of brain slices.

### Stereotaxic surgeries

Mice were anesthetized with a mixture of ketamine (100 mg/kg) and xylazine (10 mg/kg) prepared in sterile saline (intraperitoneal injection). Adequate sedation was determined by the absence of a gentle toe-pinch withdrawal reflex. To prevent ocular drying, a lubricant eye ointment (Genteal Tears; Alcon) was applied to the eyes to prevent ocular dehydration. The top of the mouse’s head was shaved, and the surgical site was disinfected using betadine. The mouse body temperature was maintained at 36.6°C by resting it on a covered heating pad (7 cm X 7 cm) connected to a rodent warmer console (Stoelting, # 53800M). To prevent somatic dehydration during surgery, each mouse was subcutaneously injected with 500 μl of sterile saline and 200μl of analgesia using 3 mg/kg subcutaneous ketoprofen (Zoetis) before surgery. Stereotaxic surgeries were carried out inside a biosafety cabinet Class II (LabRepCo, # NU-677–500) on the Kopf stereotaxic instrument (Model # 942), which was equipped with a Nanoinjector UMP3TA (WPI). The viral vectors were bilaterally injected intracranially using a 2μl Hamilton syringe (Catalog # 1127878, Syringe Neuros) at 1 nl/s into the prelimbic cortex with coordinates [−2.00 mm anterior posterior (AP), +/− 0.25 mm mediolateral (ML), and −1.60 mm dorsoventral (DV)]. The following viral vectors were used: AAV9.CamKII0.4.Cre.SV40 (Addgene, Catalog # 105558, 1.00 X 10^13 GC/ml), AAV8.hSyn.DIO.EGFP (Addgene, Catalog #50457, 1.00 X 10^13 GC/ml), AAV9.EF1a.DIO.GFP-L10 (Addgene Catalog #98747, 1.00 X 10^13 GC/ml), AAV1.CAG.GCaMP6f (Addgene, Catalog #100836, 6.80 X 10^13 GC/ml), and AAV1.EF1a-DIO.tTA (Addgene Catalog #99121, 1.00 × 10^13 GC/ml). Before removing the needle, an interval of 12 min was allotted for viral diffusion. The incision in the scalp was closed with suture clips (Fine Science Tools #12040-03) using tissue stapler (Fine Science Tools #12028-12) upon completion of the intracranial injections. After surgery, each mouse was provided with hydrogel (Clear H2O, # 70–01-5022) in the recovery cage before returning to the animal colony.

For intracerebroventricular (ICV) infusion of Asunaprevir (ASV), a 23-gauge stainless steel guide cannula (Protech International, Catalog # C315IS-5/SPC) was implanted in the left lateral ventricle (-0.50 mm AP, -1.10 mm ML, -1.60 mm DV). Dental cement containing Radiopaque L-Powder Metabond (Catalog # S396), “B” Quickbase for C&B Metabond (Catalog # S398), “C” Universal TBB Catalyst (Catalog # S371), Stoelting dental cement powder (Catalog # F51458) was then applied around the cannula’s base to securely affix the cannula in place. Once the cement solidified, the skin was sutured using the tissue adhesive Vetbond (3M). Three weeks after the surgery, 2 μl of 100 nM ASV was infused at 0.5 μl/min using an internal cannula extending out of PE50 tubing attached to a 5-ml Hamilton syringe (Hamilton) using a Syringe Infusion Pump (Pump 11 Elite Series, Item Model # 70-4504, Harvard Apparatus). The injection cannula was kept in place for 1 min before being withdrawn after the injection.

For Optic Ferrule implants, mice were unilaterally injected with 200 nl of viral cocktail into the PL (AP: +2 mm, ML: −0.25 mm, DV: -1.6 mm) and optic ferrules fabricated from 400 μm core 0.66 numerical aperture optic fiber fixed in a 3 mm diameter metal ferrule (Doric 400/430, 0.66NA, 3 mm, MF1.25 (FLT) length) were chronically implanted above the injection site in the PL with bregma coordinates (AP: +2 mm, ML: -0.25 mm, DV: -1.5 mm) 0.10 mm above the virus injection site.

### Fiber photometry

AAV injections were first performed in the left prelimbic cortex followed by optical ferrule implants. Wildtype mice received 200 nl of AAV.CAG.GCaMP6f (diluted 1:10 in saline), unilaterally into the PL (AP −2.00 mm, ML − 0.25 mm and DV −1.60 mm). Following viral infusion, imaging fibers fabricated from 400 μm core 0.66 numerical aperture optic fiber fixed in a 3 mm diameter metal ferrule (Doric 400/430, 0.66NA, 3 mm, MF1.25 (FLT) length) were chronically implanted above the injection site in the PL (AP −2.00 mm, ML − 0.25 mm, and DV −1.50 mm). Optical fibers were fixed to the skull using metabond and dental cement (Radiopaque L-Powder Metabond (Catalog # S396), “B” Quickbase for C&B Metabond (Catalog # S398), “C” Universal TBB Catalyst (Catalog # S371), Stoelting dental cement powder (Catalog # F51458)). After the ferrule implant, all mice were returned to their home cage for three weeks prior to the start of fiber photometry recording to allow for adequate viral gene expression.

Prior to performing the fiber photometry imaging experiments, all mice were habituated to patch cord tethering for 15 min. During PTC training, calcium signals were continuously monitored to ensure the acquisition of robust and reproducible data prior to the commencement of the PTC experiments. The mice were connected to an imaging patch cord (400 μm core; 0.48 numerical aperture, Doric Lenses) linked to a 6-port fluorescence minicube (Doric Lenses). Two wavelengths of light - violet (405 nm for control artifact fluorescence, Doric Lenses) and blue (465 nm for GCaMP6f excitation, Doric Lenses) - were transmitted into the brain at an intensity of 20-40 μW, which was maintained consistently across experimental sessions. The emitted light was directed through a dichroic mirror and a 500–540 nm filter before being detected by the built-in photosensor in RZ10X system. Analog signals were recorded using the RZ10X processor and a PC equipped with Synapse software (Tucker-Davis Technologies). For paired fear conditioning experiments, mice were tethered to a patch cord in context A and exposed to two CS tones (5kHz, 80 dB, 30 s) paired with two co-terminating 2 US (0.4 mA, 2 s) during the PTC training session. Following the fear conditioning experiments, the mice were returned to their home cages. Calcium signals were continuously recorded during the PTC training session and the long-term cued memory retrieval experiments, where only the CS’s were presented in Context B or C. Each photometry session commenced with a one-minute baseline period, during which calcium signals were recorded prior to the initiation of the experiment. The initiation and conclusion of each experimental session, as well as the onset and offset timestamps of the CS and US, were generated using TTL signals triggered by Graphic State software, facilitating precise temporal analysis of calcium signals.

### Translating ribosome affinity purification Sequencing (TRAP-Seq)

TRAP-Seq was performed according to the previously published work^26^. Briefly, all mice were quickly decapitated, and the PL region of interest was extracted into an ice-cold dissection buffer for washing. The collected tissues were transferred to a pre-chilled homogenizer on ice containing tissue-lysis buffer and were homogenized in a motor-driven Teflon-glass homogenizer overhead stirrer at 900 rpm with 12 complete strokes (Dounce homogenization) in 20 mM HEPES-KOH, pH 7.3, 150 mM KCl, 10 mM MgCl_2_, 1% NP-40, 0.5 mM DTT, 100 mg/ml cycloheximide, and 10 μl/ml RNasin (Promega Catalog# N2111) and 10 μl/ml SUPERase-In (Fisher Scientific Catalog # AM2696) RNase inhibitors on ice. The samples were transferred to chilled Eppendorf tubes, and nuclei were pelleted by centrifuging the lysates at 4°C for 10 min at 2,000 × g. NP-40 (Sigma; final concentration = 1%) and 07:0 DHPC (Avanti Polar Lipids Catalog # 850306P, final concentration = 30 mM) were added to the supernatant. To pellet insoluble membranes, the lysates were centrifuged at 20,000 × g, and the supernatant was transferred to a new chilled low-binding Eppendorf tube. A small aliquot of the supernatant was saved for RNA-Seq (input). A volume of 120 µl of 1 mg/mL biotinylated-Protein L (Fisher Scientific Catalog #29997) was used to coat 200 µl Streptavidin MyOne T1 dynabeads (Fisher Scientific Catalog #65601). This mixture was incubated for 35 minutes at room temperature with gentle rotation, followed by an overnight incubation at 4°C with gentle end-over-end rotation. Post-incubation, the conjugated protein L-beads were washed with 1X PBS and collected using magnets three times. The conjugated protein L-beads were then resuspended in 175 µL of 0.15 M KCl IP wash buffer (comprising 20 mM HEPES, 150 mM KCl, 5 mM MgCl_2_, 0.5 mM DTT, 1% NP-40, 100 µL RNasin Ribonuclease Inhibitor, and 100 µL SUPERase In RNase inhibitor, 100 µg/mL cycloheximide) and incubated for 1 hour at room temperature with 50 µg of each antibody. Subsequently, the polysome-bound magnetic beads were washed four times in 1,000 µL of high-salt buffer (20 mM HEPES-KOH, pH 7.3, 350 mM KCl, 10 mM MgCl2, 1% NP-40, 0.5 mM DTT, and 100 mg/mL cycloheximide). Following the fourth wash with a high-salt buffer, all residual wash buffers were removed, and the tubes were brought to room temperature. The beads were resuspended in 100 µL of Nanoprep lysis buffer with beta-mercaptoethanol (β-ME), vortexed, and incubated for 10 minutes at room temperature. The RNA was extracted from the beads using a magnet and immediately subjected to RNA cleanup according to the manufacturer’s instructions. Bound RNA was eluted and purified using the Absolutely RNA Nanoprep kit (Agilent Catalog #400753) in accordance with the manufacturer’s instructions.

The integrity of the RNA was assessed with a High Sensitivity ScreenTape (Agilent Catalog #5067-5579), and only samples with an RIN greater than 9 were selected for further analysis. RNA-Seq libraries were prepared from 50–200 pg of total RNA using the SMART-Seq v4 Ultra Low Input RNA Kit for Illumina Sequencing, with the Low Input Library Prep kit v2 (Clontech, Catalog # 634890 and 634899, respectively). Libraries were sequenced using an Illumina HiSeq 2500 instrument with a paired-end 50 protocol.

### Behavior

Before behavior testing, all animals were habituated to human experimenters for at least three days.

#### i. Open Field

Mice were placed in the center of a white plexiglas open field (27.31 cm x 27.31 cm x 20.32 cm) for 15 min. The arena was divided into center zone (inner square: 13.67 cm X 13.67 cm) and total zone encompassing the whole arena of open field. Behavioral activity was recorded using a ceiling-mounted USB camera (Logitech) and analyzed using Ethovision XT15 video tracking software (Noldus) for center body point. The parameters measured were distance traveled, and the ratio of distance traveled in the center to that in the total zone. Further, the data was analyzed as distance traveled across three time-bins, 5 min each.

#### ii. Pavlovian threat conditioning (PTC) paradigm

All animals were initially acclimated to the behavior room for 45 min prior to testing. To blunt any environmental noise, a white noise machine (Yogasleep Dohm) was used in the behavior room. Habitest Linc system - H01-01 Power Base and H02-08 Linc Boxes (Harvard Apparatus) were used in combination with the Graphic State 4 software (Coulbourn Instruments) to control delivery of footshock, light stimulus and auditory tone. Three different contexts (Context A, B and C) were used for PTC. Context A was a chamber (29 cm (W) × 23 cm (D) × 20 cm (H)) with white house light, metal shock floor, 30% ethanol, and no striped wall panels. Context B was a chamber of similar dimensions but provided with infrared light, striped wall panels, white plexiglass platform, vanilla-scented paper bedding, and 30% isopropanol. Context C was a chamber of similar dimensions in a different room and provided with red house light, blue bubble-textured rubber mat, 30% isopropanol supplemented with 100 μL peppermint oil and black wall panels. On day 1 of habituation, the mice were placed in Context A for 15 min to freely explore the chamber. For PTC training on day 2, each mouse was placed in Context A for 9.5 min to receive two pairings of conditional stimulus (CS) and unconditional stimulus (US), where CS is 5 kHz 80 dB pure tone and US is 0.4 mA footshock. The order and timing of CS and US delivery were Tone1:270s - 300s, US1:298s - 300s, Tone2:420s - 450s, US2:448s - 450s. For longitudinal retrieval groups, memory retention was tested in the same subjects on days 1 (recent LTM) and 28 (remote LTM) following training in Context B for most experiments, or in Context B and C respectively (Extended Data Fig. 1). For non-longitudinal LTM testing, mice were only tested on day 28 after PTC training in Context B. During LTM, each mouse received three CS (Tone1:270s - 300s, Tone2:440s - 470s, Tone3:570s - 600s) for 12 min. FreezeFrame 4 software (Actimetrics, version 5) was used in the recording mode to record videos and to automatically measure the freezing behavior of mice during PTC training and retrieval. Control mice in the Box Only group, that were used as controls for translation profiling or immunohistochemistry post-training, were exposed to Context A without CS and US for equivalent time as the PTC Trained mice.

### Immunohistochemistry

Mice were deeply anesthetized with isoflurane inside a fume hood and transcardially perfused with 0.1M PBS, followed by 4% paraformaldehyde (PFA; Electron Microscopy Sciences) in 1X PBS. Brains were removed and post-fixed in 4% PFA for 24 h at 4°C in a cold room on a shaker. The 4% PFA was then replaced with 30% sucrose solution for another 24 h at 4°C. All perfused brains were coronally sectioned to a thickness of 40 μm using a Leica vibratome (VT1000s) and stored in 1X PBS containing 0.05% sodium azide at 4°C. Sections were blocked with 5% normal goat serum in 0.1 M PBS with 0.1% Triton X-100 for one hour and probed overnight with primary antibodies [Chicken anti-EGFP (Abcam #ab13970, 1:1000), Guinea pig anti-cFos (Synaptic System #226004, 1:1000), and Mouse anti-eIF4E (BD Biosciences #BDB610269 1:500)]. After rinsing three times in 0.1M PBS, sliced tissue sections were subsequently incubated with Alexa Fluor conjugated secondary antibodies 1:500 (Thermo Fisher, Catalog #A32931, #A21450, #A32740, #A-11077) in blocking buffer incubated for 1.5 h at room temperature. Sliced tissue sections were washed three times in PBS for 30 min and mounted using Prolong Gold antifade mounting media with or without DAPI (Fisher # P36931, # P36930). Fluorescent images were captured using confocal microscopy (Zeiss LSM800) with 10X and 20X objective lenses. Z-projected confocal images were generated using Fiji (ImageJ).

### Western Blot

Mice were euthanized by cervical dislocation. Coronal 1 mm-thick prefrontal brain slices were cut using chilled stainless-steel brain matrix (Roboz Catalog #SA-2175). Prelimbic cortex was quickly micro-dissected in chilled 1X PBS and sonicated in ice-cold homogenization buffer (10 mM HEPES, 150 mM NaCl, 50 mM NaF, 1 mM EDTA, 1 mM EGTA, 10 mM Na4P2O7, 1% Triton X-100, 0.1% SDS and 10% glycerol) that was freshly supplemented with 10 μl each of Halt protease inhibitor (Fisher Scientific, #PI78425) and phosphatase inhibitors (Sigma # P5726, #P0044) per ml of homogenization buffer. Protein concentrations were measured using BCA assay (Fisher Scientific #23227). Samples were prepared with 5X sample buffer (0.25 M Tris-HCl pH6.8, 10% SDS, 0.05% bromophenol blue, 50% glycerol and 25% - β mercaptoethanol) and heat denatured at 95°C for 5 min. 40 μg protein per lane was run in pre-cast 4-12% NuPAGE Bis-Tris gels (Fisher Scientific #NP0322BOX) and subjected to SDS-PAGE followed by wet gel transfer to PVDF membranes (Thermo Scientific #88520). Following transfer, blots were stained for total protein using MEMCODE Reversible Protein Stain Kit for PVDF membranes (Pierce # 24585). After erasing the MEMCODE stain, blots were blocked in 5% non-fat dry milk in 0.1M PBS with 0.1% Tween-20 (PBST), and probed over 72h at 4°C using primary antibodies (rabbit anti-p-eIF2α S51 (Cell Signaling #3398, 1:300), and rabbit anti-t-eIF2α (Cell Signaling #9722, 1:500). After washing 3 times in 0.1% PBST, membranes were probed with horseradish peroxidase-conjugated secondary IgG (1:5000) (Promega # W4021) for 1h at RT. Signals from membranes were detected with ECL chemiluminescence (Pierce #PI32209) using Fluorochem E Protein Simple instrument.

### Data analysis

#### i. Fiber Photometry data analysis

To preprocess and analyze fiber photometry recordings, we implemented a multi-step artifact removal and signal normalization pipeline. Raw TDT data were first imported using the Python library tdt via a custom load_data function, which reads the .tev and .tsq files automatically generated in the specified BLOCK_PATH. This function allows selection of time windows tailored to the experimental paradigm; in our study, for training data, we extracted signals from data.epocs.PrtB.onset[0] to data.epocs.End_.offset[0], whereas for test data, we used data.epocs.PrtB.onset[0] to data.epocs.End_.onset[0]. These start and end points were empirically chosen to capture the relevant behavioral epochs while excluding extraneous signals. Consistent with recommendations from prior work (e.g., GuPPy^55^), we also removed the first 1 s of each trace to eliminate potential outlier transients at recording onset. Following data import, we applied a Butterworth low-pass filter to both the signal (465 nm) and control (405 nm) channels to suppress high-frequency noise and artifacts that can obscure the slower, biologically meaningful fluorescence dynamics associated with neural activity. Because fiber photometry signals are susceptible to gradual fluorescence decay, we performed photobleaching correction by fitting a double exponential decay function to each channel and subtracting the resulting fit from the raw traces. This step effectively removed the slow baseline drift caused by progressive fluorophore bleaching, preserving the fidelity of stimulus-evoked responses. To further address motion artifacts, which can arise from subtle animal movements during behavioral tasks, we applied a linear fit subtraction strategy: the linear relationship between the control (405 nm, isosbestic) and signal (465 nm, calcium-dependent) channels was modeled by ordinary least squares regression, and the best-fit line was subtracted from the raw signal to yield a motion-corrected trace. This approach, widely adopted in fiber photometry preprocessing, minimizes non-neural movement-induced fluctuations. After motion correction, we computed ΔF/F, defined as the percentage change in fluorescence relative to a baseline fluorescence value (𝛥𝐹/𝐹 = (𝐹 − 𝐹_𝑏𝑎𝑠𝑒𝑙𝑖𝑛𝑒_)/𝐹_𝑏𝑎𝑠𝑒𝑙𝑖𝑛𝑒_), where 𝐹 represents the motion-corrected fluorescence signal.

To enable comparisons across sessions and animals, the ΔF/F values were then converted to z-scores using the transformation 𝑧 = (𝑥 − 𝜇)/𝜎, where 𝑥 is each ΔF/F value, 𝜇 is the mean, and 𝜎 is the standard deviation of the ΔF/F distribution within the analyzed window. A positive z-score thus reflects activity above the mean, whereas a negative score indicates suppressed fluorescence. To smooth the resulting normalized signal while retaining key features such as peaks and stimulus-related transients, we applied a Savitzky–Golay (Savgol) filter with a 5000-frame window. This filter fits local polynomials over a sliding window (half-width 𝑚 such that window size = 2𝑚 + 1), generating convolution coefficients (𝑐_𝑗_) that minimize squared error and reduce noise without distorting the true underlying signal. This step is particularly advantageous for long-duration experiments, as it suppresses frame-to-frame variability while preserving biologically relevant temporal dynamics, thereby improving both visualization and downstream quantitative analyses. Finally, we computed peri-event time histograms (PETHs) aligned to the onset of the conditioned stimulus (CS) to quantify stimulus-evoked neural responses. Each PETH was generated using z-scored fluorescence traces extracted over a 90 s window (30 s before CS onset and 60 s after). For each animal, responses to individual CS presentations were calculated separately: during training, data were split into Training Instance 1 and Training Instance 2 to capture potential differences between early and later acquisition; during long-term memory (LTM) retrieval, we analyzed three CS onset events per session and averaged them to obtain daily mean responses for LTM day 1 (recent LTM), LTM day 14 (intermediate LTM), and LTM day 28 (remote LTM). The z-scored responses across instances were averaged at each time point for each day, and the resulting mean trace was plotted with the standard error of the mean (SEM) to visualize inter-trial variability. For group-level analyses, individual animal PETHs were averaged within predefined experimental groups, enabling robust comparisons across genotypes or treatments. This pipeline integrates artifact removal (motion, photobleaching, high-frequency noise), normalization (ΔF/F and z-scoring), and advanced smoothing (Savgol filtering) to yield high-fidelity, reproducible representations of neural calcium dynamics suitable for statistical and visualization purposes.

#### ii. Behavior data analysis for Pavlovian threat conditioning

To facilitate efficient and reproducible analysis of avoidance conditioning experiments, we developed a fully automated, Python-based computational pipeline for data extraction, behavioral metric computation, and visualization. The workflow integrates timestamped event logs from FreezeFrame (freezing during Pavlovian threat conditioning and signaled active avoidance) and GraphicState (conditioned stimulus [CS] and unconditioned stimulus [US] -evoked responses) to generate structured, analysis-ready datasets. Custom modules (freezeframe.py, graphicstate.py) implement preprocessing steps including column name standardization, epoch segmentation (pre-CS, CS, intertrial interval, post-CS), metadata integration, and handling of missing data. Task-specific extensions (ptc_ff.py) were generated to quantify freezing percentages across defined epochs, classify avoidance trial outcomes (avoidance, escape, failure), compute spontaneous and CS-evoked shuttling, and calculate latency to avoid using precisely parsed CS onset/offset timestamps. The pipeline uses an object-oriented modular design built around the FreezeFrame and GraphicState base classes, supporting reusability across experimental paradigms. Automated batch processing was implemented for high-throughput analysis of multiple datasets with minimal manual input. Comprehensive logging and error handling (via logging.info(), logging.warning(), try-except blocks) were also utilized to ensure data integrity and traceability. Dedicated plotting modules (fr_pav_heatmaps.py) were developed to generate standardized visualizations of freezing dynamics, allowing rapid and reproducible figure generation. Together, this custom pipeline provided a scalable and transparent framework for behavioral data analysis in Pavlovian threat conditioning paradigms.

#### iii. TRAP-seq data analysis

Raw sequencing reads (FASTQ files) were processed using nf-core/rnaseq v3.21.0dev (doi: 10.5281/zenodo.1400710), a community-curated pipeline from the nf-core collection^56^. The workflow was executed under Nextflow v24.10.6^57^, ensuring full reproducibility and portability through containerized software environments provided by Bioconda^58^ and BioContainers^59^. Reads were pre-processed with fastp for adapter and quality trimming. Contaminant sequences were screened using Kraken2/Bracken against the standard Kraken2 reference database (release 2024-09-04). For alignment and quantification, the STAR–Salmon hybrid mode was employed, mapping reads to the mouse reference genome (GRCm39, NCBI RefSeq annotation) while simultaneously generating transcript- and gene-level quantifications. Both BAM alignment files and quantification tables were retained for downstream analyses.

Principal component analysis (PCA) was performed on gene-level counts summarized with tximport and modeled in DESeq2 after filtering genes with fewer than 20 counts in at least half of the samples. Variance-stabilized expression values were computed with VST, and the top 500 most variable genes were selected for PCA using prcomp. Percent variance explained by PC1 and PC2 or PC3 was reported. Differential expression analysis was carried out in DESeq2. Genes were considered differentially expressed at adj p < 0.05 and |log2 fold change| > 0.58, with baseMean > 10 used for heatmap selection. Volcano plots displayed −log10(p or adj p) versus log2 fold change with dashed thresholds and optional y-axis breaks for extreme values, while heatmaps were generated from the top 15 up- and downregulated genes using VST. Outputs included ranked DEG tables, volcano plots, and clustered heatmaps.

Gene Ontology (GO) analysis was performed using the clusterProfiler package with org.Mm.eg.db annotations. For each contrast, genes passing significance thresholds (either *p* < 0.05 or adj p < 0.05, combined with |log₂FC| > 0.58) were separated into direction-specific sets (up in BO vs up in Trained). Gene symbols were mapped to Entrez IDs, and enrichment was run separately for Biological Process (BP), Molecular Function (MF), and Cellular Component (CC) categories. Enrichment used Bonferroni correction when based on adjusted p-values, or unadjusted when raw p-values were applied. Results were exported as tables, and the top enriched terms per direction were visualized with bubble plots showing gene counts and adjusted significance.

Rank-Rank Hypergeometric Overlap (RRHO) analysis was performed with RRHO2 to quantify threshold-free concordance between ranked gene signatures. For each contrast, genes were converted to a continuous signed score, sign(log2FC) × −log10(p_adj) (falling back to nominal p where adjusted values were unavailable), deduplicated by maximal absolute score, and ordered descending. Ranked lists were restricted to their intersecting gene set and supplied to RRHO2_initialize with log10.ind=TRUE to compute the rank–rank hypergeometric overlap surface and quadrant statistics (concordant genes: up–up, down–down, and discordant genes: up–down, down–up). Visualization comprised RRHO2 heatmaps and quadrant-specific Venn diagrams; RRHO2 objects and quadrant gene lists (list1, list2, and overlaps) were exported (RDS/CSV) under analysis-specific labels to ensure reproducibility. Cytoscape networks (TOM ≥ 0.10, top K = 400 edges/module) and module-level igraph visualizations were generated, showing the top 40 nodes (by kWithin) with the 10 highest hubs labeled.

#### iv. Image acquisition and analysis

For quantitative immunohistochemistry experiments, all imaging data were acquired using a Zeiss confocal microscope with a 20X objective lens (with 1X zoom) and z-stacks (approximately four optical sections with a 0.563 μm step size) for three coronal sections per mouse from AP Bregma +1.98 mm to + 1.50 mm (n = 3 mice) were collected. For assessing viral targeting, the imaging data were acquired using a 10X objective lens (with 1X zoom) and tiled scan feature spanning the hemisphere of the coronal prefrontal brain section. ImageJ was used to analyze the imaging data using the Bio-Formats importer plugin. Initially, a maximum projection of the z-stacks was generated, followed by manual outlining of individual cells to measure their mean fluorescence intensity using drawing and measurement tools. Background fluorescence of 5 ROIs equivalent in size to the cell ROIs was measured and subtracted from the mean intensity. Fluorescence intensity values from all cell measurements were further normalized to the mean fluorescence intensity of the control samples.

#### v. Immunoblot densitometry – image analysis

Images of immunoblots following western blotting were opened in Fiji and bands were outlined and selected using the Analyze: Gels plugin – Select First Lane and Select Next Lane features. Plot Lanes feature was used to generate a trace of the band density, and Area under the curve was selected manually using a line drawing tool followed by intensity measurement using the wand tool. The mean intensity measures were normalized by total protein and averaged across multiple replicates.

### Statistics

Statistical analyses were performed using GraphPad Prism 9 (GraphPad software, version 9.5.1). Data are expressed as mean ± SEM for all X-Y plots, and as Median ± Interquartile for all violin plots. Data from two independent groups were analyzed using a two-tailed unpaired Student’s t-test. Multiple group comparisons were performed using analysis of variance (ANOVA), followed by appropriate post hoc tests, as described in the figure legend. Statistical significance was set at p < 0.05.

